# Amyloid β oligomers constrict human capillaries in Alzheimer’s disease via signalling to pericytes

**DOI:** 10.1101/357095

**Authors:** Ross Nortley, Anusha Mishra, Zane Jaunmuktane, Vasiliki Kyrargyri, Christian Madry, Hui Gong, Angela Richard-Loendt, Sebastian Brandner, Huma Sethi, David Attwell

## Abstract

Vascular compromise occurs early in Alzheimer’s disease (AD) and other dementias^1–3^. Amyloid β (Aβ) reduces cerebral blood flow^4–6^ and, as most of the cerebral vasculature resistance is in capillaries^7^, Aβ might mainly act on contractile pericytes on capillary walls^8–10^. Employing human tissue to establish disease-relevance, and rodent experiments to define mechanism, we now show that Aβ constricts brain capillaries at pericyte locations in human subjects with cognitive decline. Applying soluble Aβ_1-42_ oligomers to live human cortical tissue constricted capillaries. Using rat cortical slices, this was shown to reflect Aβ evoking capillary pericyte contraction, with an EC_50_ of 4.7 nM, via the generation of reactive oxygen species and activation of endothelin ET-A receptors. In freshly-fixed diagnostic biopsies from human patients investigated for cognitive decline, mean capillary diameters were less in subjects showing Aβ deposition than in subjects without Aβ deposition. For patients with Aβ deposition, the capillary diameter was 31% less at pericyte somata than away from somata, predicting a halving of blood flow. Constriction of capillaries by Aβ will contribute to the energy lack^1–3^ occurring in AD, which promotes further Aβ generation^11,12^. This mechanism reconciles the amyloid hypothesis^13–15^ with the earliest events in AD being vascular^1^.

---

Around the time that the amyloid hypothesis for Alzheimer’s Disease (AD) was published^13–15^, it was reported that capillaries in the brains of AD patients showed an abnormal focally-constricted morphology^16,17^ similar to that produced by pericyte contraction^8,9^. Although much succeeding work focused on Aβ- and tau-evoked damage to neurons, increasing evidence suggests a role for vascular disturbance in the onset of AD: an idea formalised in 1993^18,19^. This concept is supported by observations of reduced cerebral blood flow early in the development of the disease^1^, and by the fact that reduced cerebral blood flow increases Aβ production^11,12^.

Investigations of the vascular effects of Aβ and AD have focused on arteries and arterioles^5,6,20^, but in the CNS the majority of the vascular resistance is located in capillaries^7^. A subset of pericytes on capillary walls is contractile and can alter cerebral blood flow by adjusting their contractile tone^8,9^, and in a rodent model of AD there are disturbances of unknown origin in the control of capillary blood flow^10^. We therefore investigated how pericytes were affected by Aβ_1–42_ oligomers, the molecular species believed to be responsible for Aβ’s toxic effects in AD^21,22^. To maximise the relevance to human disease, unlike most AD-related studies on rodents, we used living human brain slices derived from neurosurgically-resected brain tissue (removed to access tumours) to study acute responses to Aβ, and rapidly-fixed human brain biopsy tissue (with or without Aβ deposition) to assess pericyte responses to long-term accumulation of Aβ in AD. The mechanisms of the effects seen in human tissue were defined in rodent brain slices.

Living human brain cortex slices were obtained from tissue removed during neurosurgical operations to access tumours (see Online Methods) and either fixed for immunohistochemistry or imaged live to study pericyte properties. Labelling the basement membrane with fluorescently-tagged isolectin B_4_, or immunolabelling for the pericyte marker platelet-derived growth factor receptor (PDGFRβ), revealed pericyte morphology. Pericytes were observed with a classical “bump-on-a-log” morphology on the straight parts of capillaries (Fig. 1a), or at their branch points, with processes extending along and around the capillaries (Fig. 1b). The mean distance between pericytes was 65.3±0.4 μm (for 94 pericytes imaged in tissue from 2 patients), 30% larger than in rodents^9^. As for arteriole smooth muscle (Fig. 1c), the processes of 36% of pericytes labelled for α smooth muscle actin (Fig. 1d), providing a mechanistic substrate for the Aβ-evoked contraction described below. Superfused noradrenaline constricted and glutamate dilated the capillaries at pericyte locations (Fig. 1e, f), as previously reported for rodent capillaries^8^, confirming that the surgery-derived human tissue had functioning contractile pericytes.

**Figure 1.**
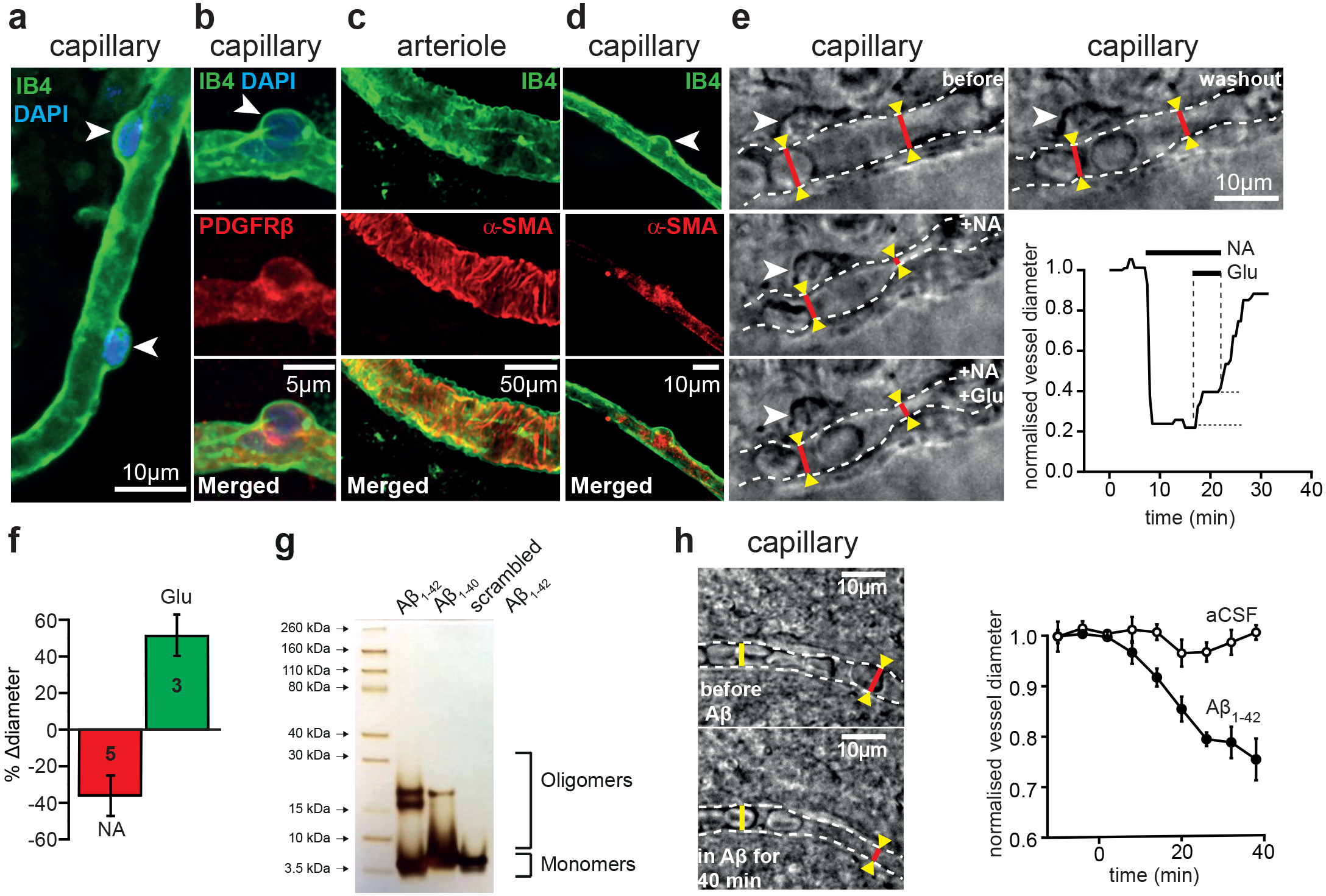
Oligomeric Aβ acts on pericytes to constrict capillaries in human brain slices. (**a**) Isolectin B_4_ labelled capillary in a human cortical slice, with two pericyte somata (arrowheads) outlined by their basement membrane. (**b**) Pericyte labelled with antibody to PDGFRβ. (**c-d**) Arteriole (**c**) and pericyte (**d**) labelled with isolectin B_4_ and antibody to α smooth muscle actin (SMA). (**e**) Images of a capillary (dashed lines indicate wall) and pericyte soma (arrowhead) in a live brain slice before drug application (*before*), in the presence of 2 μM superfused noradrenaline (+NA), with 2 μM NA and 500 μM glutamate superfused (+NA +Glu), and after stopping drug superfusion (*washout*). Graph shows time course of capillary diameter at right red line throughout the experiment. (**f**) Mean glutamate-evoked dilation and noradrenaline-evoked constriction in experiments as in (e) (change in diameter quantified relative to that before application of each drug; relative to the pre-noradrenaline diameter the glutamate-evoked dilation was 26.8±7.7%). (**g**) Silver staining of an SDS-PAGE gel for Aβ solutions prepared as in the Online Methods. (**h**) Images of a capillary before and after superfusion of 72 nM Aβ_1–42_, showing a region (red line) being constricted by pericytes and one region (yellow line) that is not constricted. Graph shows mean diameter change at 4 pericyte locations from 3 slices treated with Aβ and 3 pericyte locations from 3 slices treated with aCSF (significantly reduced at 40 mins in Aβ, p=0.01).

Aβ was oligomerised (see Online Methods), and silver staining of SDS-PAGE gels was used to assess the degree of aggregation of the Aβ isoforms. The predominant species produced (other than monomers) for Aβ_1-42_ and Aβ_1-40_ had a molecular weight of 4 to 5 times that of monomers, whereas scrambled Aβ_1-42_ formed mainly monomers (Fig. 1g). Applying soluble Aβ_1-42_ (oligomeric + monomeric, 72 nM) to human brain slices evoked a slowly developing constriction of capillaries that reduced their diameter by ~25% (p=0.01) after 40 mins (Fig. 1h). For a fixed pressure across the capillary bed, a focal constriction of this magnitude at pericyte somata is predicted to reduce cerebral blood flow by 45% if it occurred throughout the capillary network (see Online Methods).

As the limited availability of live human tissue precludes detailed analysis of the mechanism underlying the Aβ-evoked constriction, we carried out experiments on cortical slices from 21-day-old rats to investigate this. As for human capillaries, Aβ_1-42_ and also Aβ_1-40_ evoked a constriction of rat capillaries near pericyte locations that was visible using either bright field illumination or 2-photon fluorescence imaging of isolectin B_4_ (Fig. 2a-c). The time course of the Ap-evoked constriction (Fig. 2c) was similar to that in human cortex (Fig. 1h), reaching ~16% after 1 hour (p=0.006 for Aβ_1-42_ and 0.048 for Aβ_1-40_). Capillaries monitored for an hour without applying Aβ, or those to which a version of Aβ_1-42_ with a scrambled sequence was applied (prepared as for the Aβ oligomers), showed no significant diameter change (Fig. 2c). Scrambled Aβ_1-42_ does not form oligomers (Fig. 1g), unlike Aβ_1-42_ and Aβ_1-40_, which may indicate that oligomer formation is obligatory for an effect on pericytes. The pericyte-mediated constriction evoked by Aβ_1-42_ showed a Michaelis-Menten dependence on Aβ concentration, with an apparent EC_50_ of 4.7 nM (Fig. 2d).

**Figure 2.**
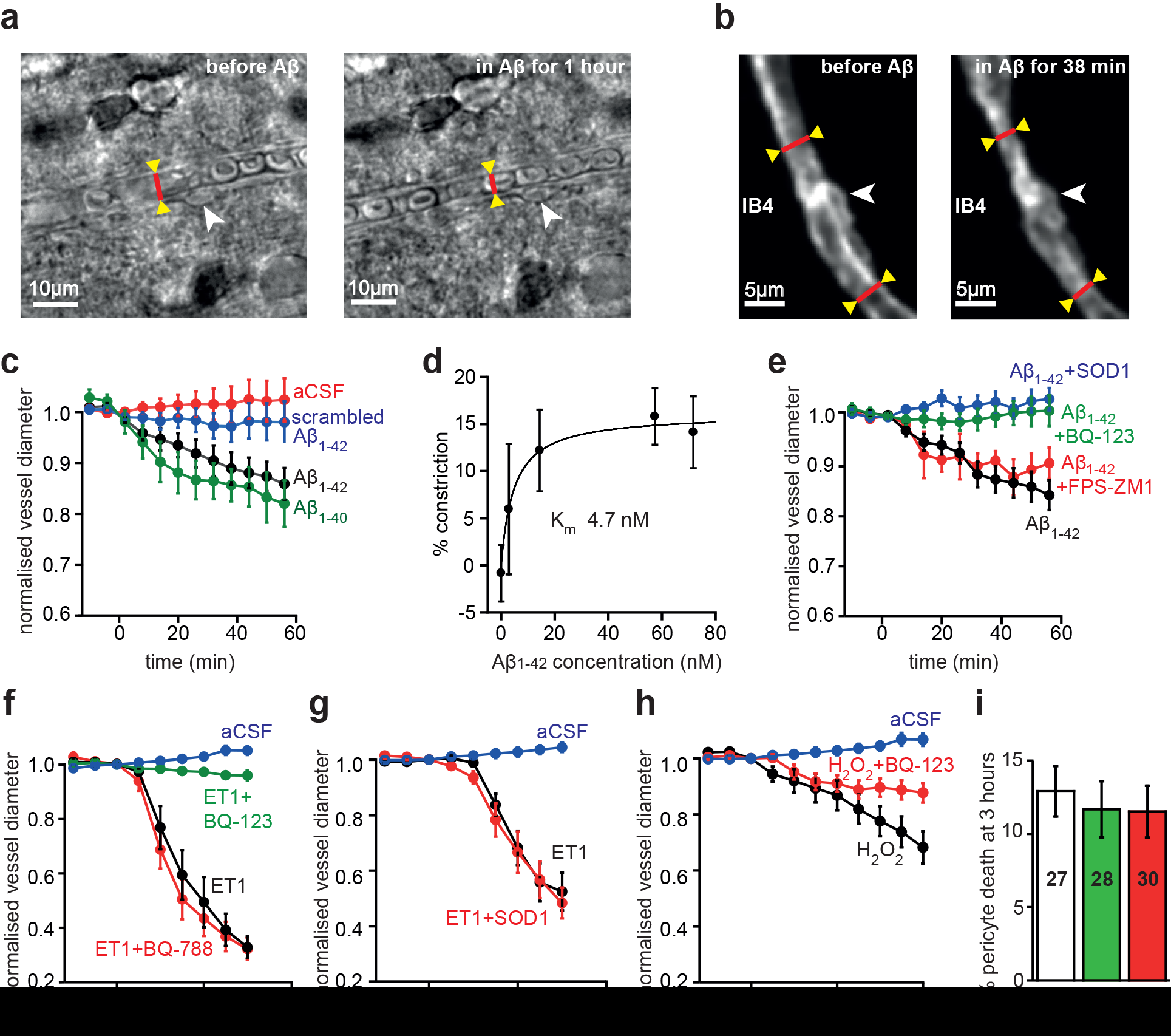
Aβ acts via reactive oxygen species and endothelin type A receptors. (**a-b**) Bright field images (**a**) and 2-photon images of IB_4_ fluoresence (b) of capillaries in rat cortical slices in normal aCSF and after superfusion with 72 nM Aβ_1-42_, showing capillary constriction near pericytes. (**c**) Mean time course of normalised capillary diameter during superfusion with aCSF (n=51 vessels), scrambled Aβ_1-42_ (109 nM, n=32), Aβ_1-42_ (72 nM, n=20) or Aβ_1-40_ (100 nM, n=6). (**d**) Constriction produced after 1 hour by different concentrations of Aβ_1-42_ (n=51, 11, 10, 19 and 20 for 0, 2.9, 14, 57 and 72 nM respectively). Curve through the points is a Michaelis-Menten relation with a best-fit K_m_ of 4.7 nM and a maximum constriction of 16.1%. (**e**) Mean time course of normalised capillary diameter during superfusion with 57 nM Aβ_1-42_ alone (n=19), or in the presence of superoxide dismutase 1 (SOD1, 150 units/ml, n=19), the ET-A receptor blocker BQ123 (1 μM, n=14), or the RAGE blocker FPS-ZM1 (1 μM, n=8). (**f**) Mean time course of normalised capillary diameter during superfusion with aCSF (n=10), ET1 alone (10 nM, n=10), or ET1 in the presence of the ET-A receptor blocker BQ-123 (1 nM, n=10) or the ET-B receptor blocker BQ-788 (1 μM, n=12). (**g**) SOD1 (150 U/ml, n=8) does not block the constriction evoked by endothelin (ET1, 5 nM, n=12). (**h**) The reactive oxygen species generator H_2_O_2_ (1 mM, n=9) evokes capillary constriction (p=1.1×10^−5^ at 20 mins) which is attenuated by the ET-A receptor blocker BQ123 (1 μM, n=11, p=0.009). (**i**) Incubating rat brain slices in solution containing Aβ_1-42_ oligomers (1.4 μM, n=28) or ET1 (100 nM, n=30 slices) for 3 hours does not increase pericyte death over that seen when incubating in aCSF (n=27).

The Aβ_1-42_-evoked capillary constriction in rat cortical slices was blocked by the endothelin-1 (ET-1) type A receptor blocker BQ123 (1 μM, p=0.008), or by application of superoxide dismutase 1 (SOD1, 150 units/ml, p=3.7×10^−6^) which scavenges reactive superoxide generated when Aβ activates NADPH oxidase in immune cells (Fig. 2e), as previously reported for the effect of Aβ on arteries and isolated penetrating arterioles^5,6,20^. Neither of these agents alone affected capillary diameter (BQ123: −0.7±5.2% dilation, n=13; SOD1: 3.4±5.8% dilation, n=9). Unlike for Aβ_1-40_ effects on blood flow *in vivo*^5^, blocking binding of Aβ to the receptor for advanced glycation endproducts (RAGE) using FPS-ZM1 (1 μM) had no effect on the pericyte constriction (p=0.34, Fig. 2e), reflecting the fact that we applied Aβ in the extracellular fluid: RAGE is only needed to transport Aβ into the brain from the blood^5^. To confirm that pericytes constrict in response to activation of ET-1 type A receptors, we applied ET-1 (10 nM) either alone or with a blocker of its type A or type B receptors. ET-1 evoked a strong pericyte-mediated constriction of capillaries (p=2×10^−12^), which was blocked by the type A receptor blocker BQ123 (1 μM, p=2.6×10^−11^), but not by the type B receptor blocker BQ788 (1 μM, p=0.91, Fig. 2f). ET-1 still evoked a constriction in the presence of SOD1 (p=1.3×10^−8^, Fig. 2g) implying that ET-1 acts downstream of superoxide, while generating reactive oxygen species using H_2_O_2_ evoked a constriction (p=1.1×10^−5^) that was reduced by BQ123 (p=0.009, Fig. 2h) suggesting that H_2_O_2_ constricts pericytes via endothelin receptor activation. These data establish Aβ-evoked generation of reactive oxygen species, presumably by resident microglia^23^ or perivascular macrophages^24^, as being upstream of the elevated level^25,26^ (or potentiated effect^27^) of endothelin which makes pericytes constrict capillaries.

In profound ischaemia, pericyte-evoked constriction of capillaries is followed by the pericytes dying in rigor (probably^9^ via an excessive rise of [Ca^2+^]_i_), thus maintaining a decreased capillary diameter and a long-lasting decrease of blood flow^9^, and pericytes also die after accumulating Aβ in AD^28^. We assessed whether exposure to 1.4 μM soluble Aβ_1-42_ or 100 nM ET-1 for 3 hours had a similar effect on pericyte health, by applying propidium iodide to label cells with membranes that had become non-specifically permeable^9^. These procedures did not significantly increase pericyte death on this time scale (Fig. 2i, p=0.85 for Aβ_1-42_ and 0.59 for ET-1).

Since acute exposure to Aβ cannot mimic the slow increase which occurs over decades in human AD patients, we studied rapidly-fixed brain cortical biopsy tissue from patients being investigated for cognitive decline of unknown cause. Tissue sections were labelled with antibodies to Aβ (recognising residues 8-17 of Aβ), PDGFRβ and phosphorylated tau. Of 13 patients (for demographics, see Online Methods), 7 turned out to have Aβ deposition while 6 did not. Specimen images of their vessels and Aβ labelling are shown in Fig. 3a, b. Pericytes were readily identifiable from their PDGFRβ labelling. Averaging over 120-140 adjacent fields of view (400 μm square in size, randomly placed on each section as a 5×4 grid of squares) in tissue from the two types of patient, with the analyst blinded to the occurrence of Aβ deposition (viewing only the PDGFRβ image channel), we found that the mean capillary diameter was reduced by 8.1% (p=0.0007) in the patients with Aβ deposition (5121 diameters measured) compared to those without Aβ deposition (3921 diameters measured, Fig. 3c).

**Figure 3.**
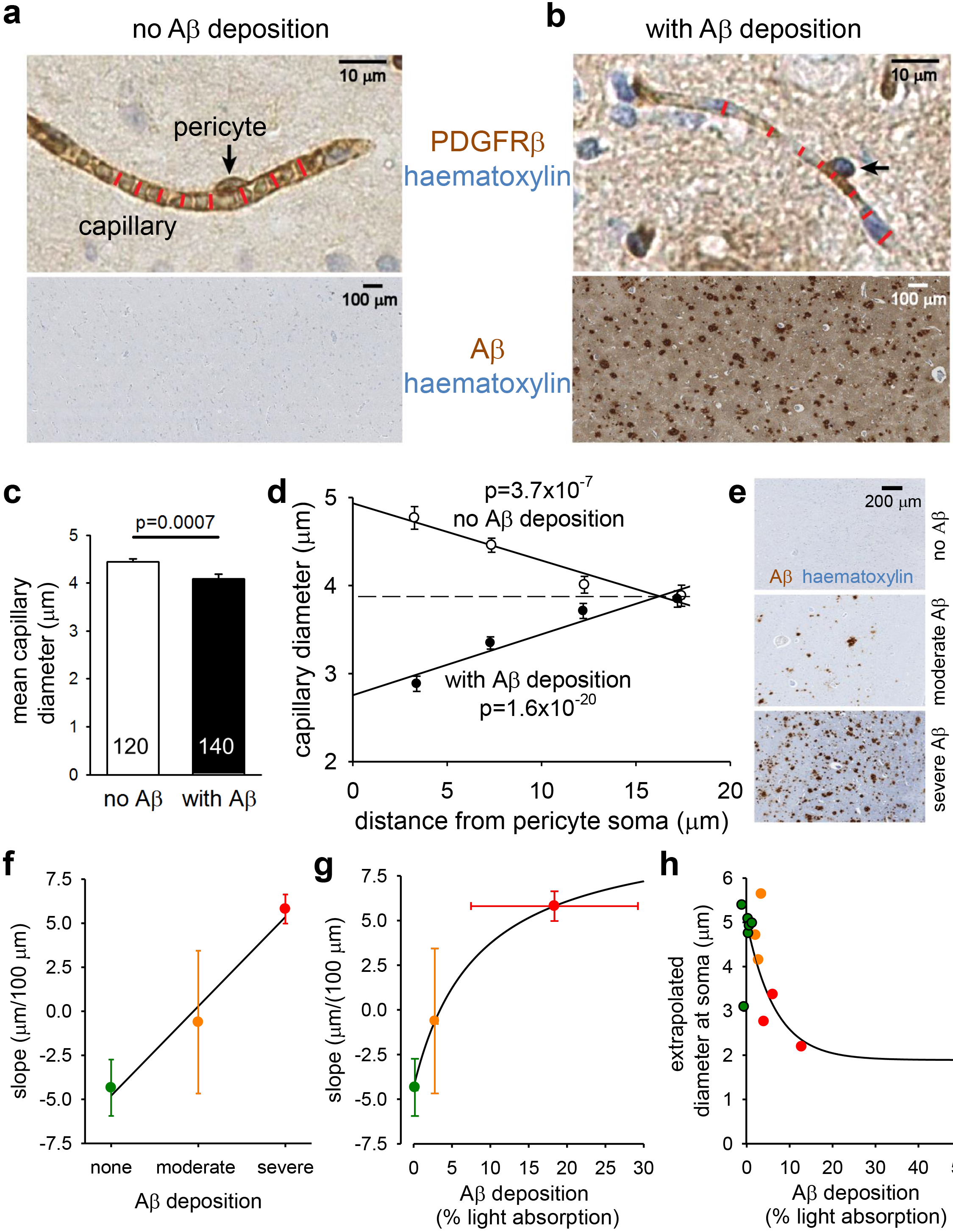
Pericyte-mediated capillary constriction occurs in humans with Aβ deposits. (**a-b**) Sample images of human cortical biopsies, labelled for PDGFRβ (brown in top panels) to show pericytes (arrows), from patients lacking (**a**) or exhibiting (**b**) Aβ deposits (brown in bottom panels, haematoxylin counterstain blue). Red lines indicate capillary diameter. (**c**) Mean diameter of capillaries in patients lacking (3921 diameters measured) or exhibiting (5121 diameters measured) Aβ deposits (number of images analysed shown on bars). (**d**) Dependence of capillary diameter on distance from a visible pericyte soma (in 5 μm bins, from 0-5, 5-10, 10-15 and 15-20 μm) for patients lacking or exhibiting Aβ deposits (moderate and severe Aβ deposition pooled together). P values assess whether slope of regression line is significantly different from zero. (**e**) Examples of Aβ labelling assessed by the neuropathologist as absent, moderate or severe. (**f**) Slope of regression lines as in d plotted as a function of neuropathologist-rated parenchymal Aβ load. (**g**) Slope of regression lines as in d plotted as a function of severity of Aβ deposition measured optically, with subjects grouped by colour (defined in f) as classified by neuropathologist. (**h**) Dependence of extrapolated diameter at soma (as in d) on severity of Aβ deposition measured optically, with subject points coloured as classified by neuropathologist (colours defined in f). Lines through data in f-h are to show the trends in the data.

To assess whether this diameter reduction was a non-specific effect of AD, or was pericyte-related, we plotted the capillary diameter measurements as a function of the distance from the nearest PDGFRβ-labelled pericyte soma (see Online Methods). In patients with no detectable Aβ deposition the capillary diameter increased at locations near pericyte somata compared to at locations far from the soma (~25% larger, slope of line is significantly less than zero, p=3.7×10^−7^ for 813 data points from 6 such patients, Fig 3d). A similar increase in capillary diameter near somata has previously been found in rodent brain capillaries *in vivo*^9^, and attributed to the presence of the soma inducing more growth of the endothelial tube. In contrast, in patients with Aβ deposition, the capillary diameter was significantly reduced near the pericyte somata compared to locations distant from the somata (Fig. 3d; ~30% smaller, slope of line is significantly greater than zero, p=1.6×10^−20^ for 1313 data points from 7 patients), as expected if pericytes cause the capillary constriction by contracting their circumferential processes which are mainly located near their somata. This constriction is predicted to reduce flow by ~50% compared to if there were no constriction (see Online Methods).

The subjects were classified by neuropathologists assessing the Aβ-labelled biopsies as having “no Aβ deposition”, “moderate Aβ deposition” or “severe Aβ deposition” in the parenchyma (as diffuse deposits and/or as plaques with central amyloid cores, Fig. 3e). The mean slope from graphs like those in Fig. 3d, for 6 patients with no Aβ deposition, 3 patients with moderate deposition and 4 patients with severe Aβ deposition, showed a progressive change from negative (implying a larger capillary diameter at the soma) to positive (implying a smaller diameter at the soma) as the severity of the Aβ deposition increased (Fig. 3f, p=0.003 compared with a relationship with zero slope), supporting further the idea that Aβ is the cause of the capillary constriction. Plotting a similar graph for the slope against the neuropathologist-assessed degree of cerebral amyloid angiopathy or phosphorylated tau labelling, gave a broadly similar trend with disease severity, but with a somewhat less significant p value (0.013 and 0.007 respectively).

To quantify Aβ level more rigorously, we measured light absorption by the peroxidase product generated by the Aβ antibody, in the region where the vessel diameters were measured in each biopsy (see Online Methods; although this measure of Aβ may largely reflect the presence of plaques, it is likely that the soluble Aβ concentration correlates with plaque load). Plotting the slopes of graphs like those in Fig. 3d, for each biopsy, as a function of the amount of Aβ deposition, again showed a monotonic progression from a negative slope to a positive slope as Aβ deposition increased, but with the change of slope occurring more strongly at low levels of Aβ deposition (Fig. 3g). Similarly, plotting the value of the capillary diameter at the pericyte soma for each biopsy (extrapolated from the straight line fit) as a function of Aβ deposition showed that the diameter was reduced more strongly by low levels of Aβ, with less further constriction as deposition increased (Fig. 3h).

Our data demonstrate that, at low nanomolar concentrations, soluble Aβ_1-42_ oligomers evoke a constriction of human cortical capillaries, mediated by pericytes. Capillaries are the site in the cortical vasculature where most of the resistance to flow is located^7^, and so are the site where diameter changes will reduce blood flow most (although arteries^20^ and arterioles^6^ are also reported to be constricted by Aβ). In rodents we show that this constriction is the result of reactive oxygen species generation activating endothelin signalling via ET_A_ receptors, which makes pericytes constrict the capillaries. The EC_50_ for Aβ’s action, 4.7 nM, is comparable to the concentration of soluble Aβ found in the brain (6 nM, from the TBS fraction in Table 1 of ref. 29) and two factors indicate that the effects seen are pathologically relevant. First, analysing the diameter of capillaries in human patients with cognitive decline, who either showed or lacked deposition of Aβ, shows that Alzheimer’s pathology leads to capillary constriction specifically at pericytes, with a magnitude that is estimated to reduce blood flow by a factor of at least **2**. Second, the magnitude of the capillary constriction in dementia patients increases with the severity of Aβ deposition.

Both the reduction of basal blood flow produced by Aβ, and a reduction in the blood flow increase normally produced by neuronal activity^30^ (which may also reflect the action of Aβ on pericytes^9^), will decrease the energy supply to the brain. This in turn increases Aβ production by upregulating β-amyloid converting enzyme (β-secretase 1, BACE1)^11,12^. Consequently, the regulation of pericyte-mediated capillary constriction by Aβ may act as an amplifying mechanism in a positive feedback loop (Fig. 4), increasing the level of Aβ and tau aggregation which ultimately lead to the loss of synapses and neurons.

**Figure 4.**
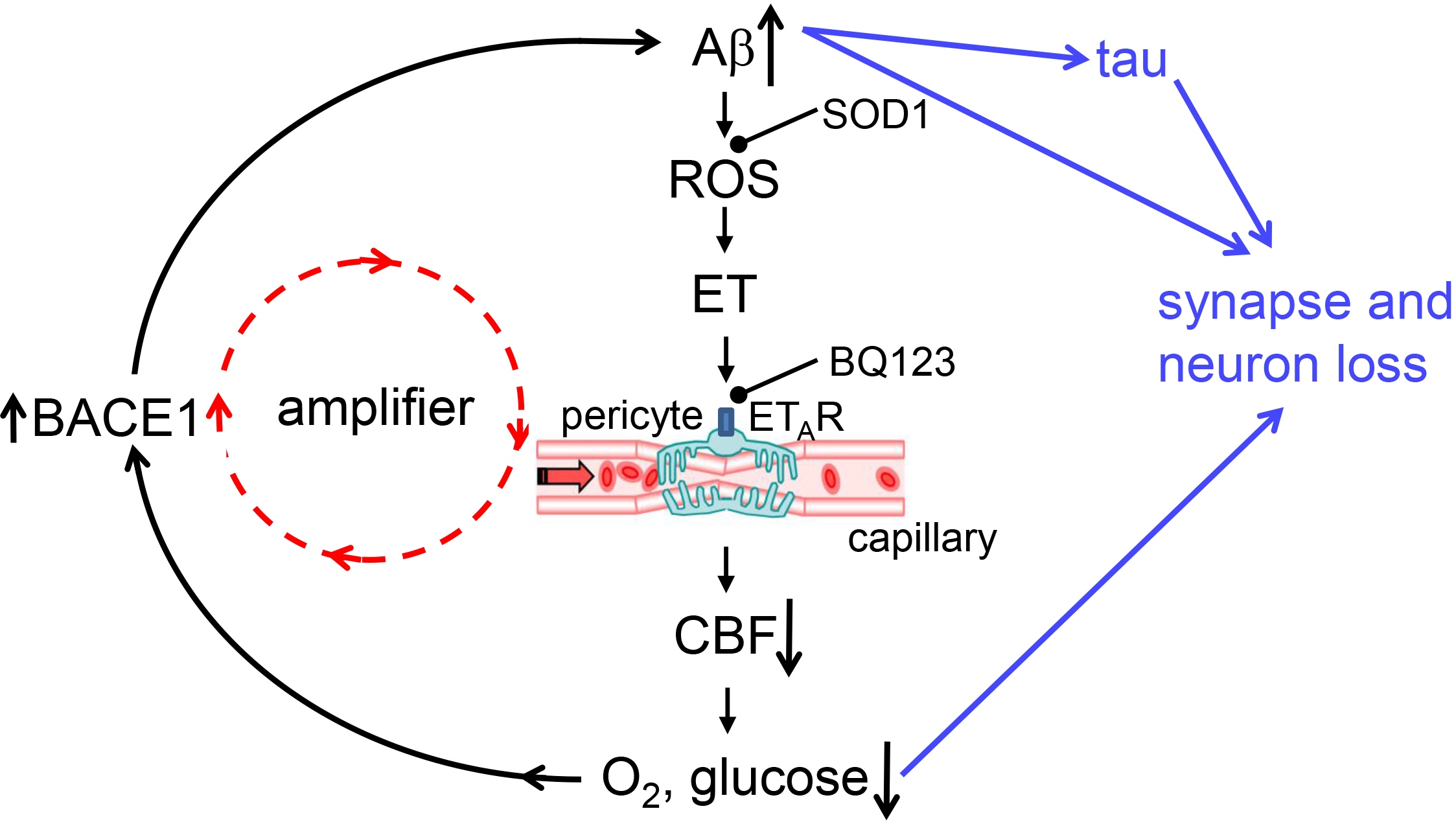
Aβ effects on capillaries may amplify the onset of AD. Amyloid β oligomers activate NADPH oxidase in immune cells^23,24^ to generate reactive oxidative species (ROS). These in turn release, or potentiate the constricting effects of, endothelin which acts via ET_A_ receptors on pericytes on capillaries - the locus^7^ of the largest component of vascular resistance within the brain parenchyma. Capillary constriction decreases cerebral blood flow and hence the supply of oxygen and glucose to the brain, which increases the production of Aβ, in part by upregulating^11,12^ expression of β-amyloid converting enzyme (β-secretase 1, BACE1), thus forming an amplifying positive feedback loop. Either directly, or via tau production, or via the decrease in oxygen and glucose supply, a rise in Aβ concentration leads to the loss of synapses and neurons. Potential sites for therapeutic intervention are highlighted at the stages of ROS (SOD1) and endothelin receptors (BQ123).

These data suggest several potential therapeutic approaches for early AD, based on the mechanisms generating pericyte constriction. Aβ-evoked generation of reactive oxygen species (ROS) might be targeted, but this would require resolving whether these are generated by resident microglia^23^ or perivascular macrophages^24^, and ROS may play important signalling roles in pathways other than the one controlling endothelin effects. A more promising approach, therefore, might be to try to reduce endothelin release (which is presumably from microglia or endothelial cells: the brain cells expressing endothelin strongly) or the effects of endothelin specifically on its type A receptors located on CNS pericytes. This could be achieved, for example, by generating an endothelin receptor blocker which also binds to another target protein on pericytes, such as the proteoglycan NG2. This concept raises the question of whether Aβ also evokes constriction of capillaries in other vascular beds by acting through pericytes, in which case drugs might be usefully targeted at all pericytes, and not just those in the brain.

## Acknowledgements

Supported by European Research Council (BrainPower and BrainEnergy) and Wellcome Trust Investigator Awards (099222/Z/12/Z) to DA, support to SB from the National Institute of Health Research (NIHR) UCLH/UCL Biomedical Research Centre, and a PhD studentship to RN from the Leonard Wolfson Experimental Neurology Centre. We thank Narges Bazargani, Bart De Strooper, Marc Ford, Nick Fox, Alastair Gibb, John Hardy, Pablo Izquierdo, Josef Kittler, Colin Masters, Patricia Salinas and Angus Silver for comments on the manuscript.

## Online Methods

### Human brain slices

The work received ethical approval from the National Health Service (REC number 15/NW/0568) and all patients gave informed consent. During neurosurgical operations for tumour treatment, apparently normal cortical tissue that was removed (to gain access to the tumour), which would otherwise have been discarded, was placed in ice cold brain slicing solution containing (mM) 93 N-methyl-D-glucamine (NMDG) chloride, 2.5 KCl, 30 NaHCO_3_, 10 MgCl_2_, 1.2 NaH_2_PO_4_, 25 glucose, 0.5 CaCl_2_, 20 HEPES, 5 Na ascorbate, 3 Na pyruvate, 1 kynurenic acid (to block glutamate receptors, to prevent excitotoxic damage to neurons), oxygenated by gassing with 95% O_2_/5% CO_2_, and transported in less than 15 mins to the laboratory. Tissue was cut into 200 μm sections and the slices were incubated at 34°C in the same solution for 10 mins, and then incubated at room temperature until used in experiments in a similar solution^31^ with the NMDG-Cl, MgCl_2_ and CaCl_2_ replaced by (mM) 92 NaCl, 1 MgCl_2_ and 2 CaCl_2_. Each patient’s tissue typically generated ~2 brain slices. When sufficient tissue was present, histological examination of the slices using haematoxylin and eosin by neuropathologists was used to assess tumour infiltration into the nominally normal tissue. This revealed that while some slices showed no infiltration by the tumour, others did. Aβ was only applied to slices that showed no tumour infiltration. Pericyte responses to noradrenaline and glutamate as documented in Fig. 1 were observed whether or not there was tumour infiltration.

### Rodent brain slices

Experiments used P21 Sprague-Dawley rats of either sex. All animal procedures were carried out in accordance with UK regulations. Cerebral cortical slices (300 μm thick) were prepared^8^ and stored as for human slices.

### Extracellular solution

Brain slices were superfused at ~4 ml/min with solution containing (mM) 124 NaCl, 2.5 KCl, 26 NaHCO_3_, 1 MgCl_2_, 1 NaH_2_PO_4_, 10 glucose, 2 CaCl_2_. This solution was gassed with 20% O_2_/75% N_2_/5% CO_2_, which produces a physiological level of oxygen in the slice near the capillaries being imaged^9^.

### Imaging capillaries

Healthy capillaries (<10 μm in diameter, mean diameter 5.61±0.03 μm (n=299) in rat, and 5.08±0.33 μm (n=12) in human, with no rings of arteriolar smooth muscle around them) were selected as previously described^31^, and regions of them were imaged which were in focus in a single image plane over at least 30 μm along the length of a capillary and exhibited a candidate pericyte with a bump-on-a-log morphology (Figs. 1e, 2a). A CCD camera was then used to capture images 100 μm square during superfusion of drugs. Capillary diameter was measured from the resulting movies, by an analyst blinded to the time and identity of drug application, by placing a line across the lumen on magnified images. In some experiments pericytes were identified prior to imaging by incubating slices for 30 min in 10 μg/ml isolectin B_4_ conjugated to Alexa 488, which binds to α-D-galactose residues in the basement membrane secreted by pericytes and endothelial cells, and outlines pericytes^31^. This allowed 2-photon imaging (using a Zeiss LSM710 microscope, excitation wavelength 800 nm) of the endothelial tube and the pericytes on it (Fig. 2b).

### Assessing pericyte death

This was carried out as previously described^9^. Briefly, brain slices (250 μm thick) were incubated in a multi-well plate, with oxygen blown gently at the surface, in aCSF, or aCSF with oligomerised Aβ_1-42_ or ET-1 added. All extracellular solutions contained isolectin B_4_ to label the basement membrane, and hence pericytes which are enveloped by this (Fig. 1b), and 7.5 μM propidium iodide to label cells with membranes that had become non-specifically permeable^9^. After 3 hours incubation, slices were fixed in 4% paraformaldehyde, and imaged on a confocal microscope. To avoid counting cells killed by the slicing procedure, quantification of the percentage of pericytes that were dead excluded cells within 20 μm of the slice surface.

### Oligomerising Aβ and assessing the form and concentration of Aβ applied

The method employed to generate oligomeric Aβ preparations was modified from that previously described^32^. Synthetic Aβ_1-42_ (Bachem H-1368.1000), Aβ_1-40_ (Bachem H-1194.1000) and scrambled Aβ_1-42_ (Bachem H-7406.1000) were suspended in 1,1,1,3,3,3 hexafluoro-2-propanol (HFIP; 52527, Sigma) at 1 mM, vortexed to obtain a homogenous solution, and aliquoted to microcentrifuge tubes. The HFIP was removed by overnight evaporation and the Aβ was completely lyophilised via a Speed-Vac. The Aβ peptide films were stored desiccated at −20°C until further processed. The peptide films were then resuspended at a nominal 5 mM in DMSO, bath-sonicated for 10 min and vortexed for 30 sec. To form Aβ oligomers, this solution was diluted to a nominal 100 μM with phosphate-buffered saline, vortexed for 15-30 sec and incubated at 4°C for 24 h. Immediately before use, the oligomeric preparations were centrifuged at 14,000 g for 10 min at 4°C (to remove any fibrils that might be present) and the supernatants were further diluted to the final experimental concentrations (quantified below) with extracellular solution.

Quantification of Aβ peptide concentration was performed using a Pierce BCA protein assay kit (Thermoscientific 23227), calibrated against a known concentration of bovine serum albumin, taking into account the different chromophoric development of albumin and Aβ peptides by multiplying by a factor of 1.5 1 ^33,34^. This showed that the amount of the molecule remaining as soluble monomers and oligomers was 28.7±2.9% (n=4) of the total concentration added for Aβ_1-42_, 39.9±1.5% (n=3) for Ap_1-40_, and 43.6±2.3% for scrambled Aβ_1-42_. Concentrations stated in the text have been corrected for these factors.

The Aβ oligomeric preparations were analysed via SDS-PAGE using 10-20% tris-glycine gels (EC61352BOX, Invitrogen). Samples of 50 μg Aβ peptides were added to tris-glycine SDS Sample Buffer (LC2676, Invitrogen). Equal volumes of each sample (10 μl) were loaded onto gels along with SeeBluePlus2 (Invitrogen) pre-stained molecular weight markers, and electrophoretically separated at 100 V. Gels were stained for total protein using a SilverXpress Silver Staining kit (LC6100, Invitrogen) according to the manufacturer’s protocol. Aβ_1-42_ and Aβ_1-40_ formed monomers and oligomers, while scrambled Aβ_1-42_ formed only monomers (Fig. 1g).

### Human biopsy data

Diagnostic brain biopsies, comprising cortex and subcortical white matter, were performed as part of routine clinical investigation at the National Hospital for Neurology and Neurosurgery, Queen Square, London, to exclude treatable causes of neurological symptoms the patients had presented with. All patients gave informed consent for the biopsy. The use of human tissue samples was licensed by the National Research Ethical Service, UK (University College London Hospitals NRES license for using human tissue samples, project ref 08/0077). The storage of human tissue was licensed by the Human Tissue Authority, UK (License #12054).

Biopsies (volume typically 1 cm^3^) were all from the right frontal lobe. The biopsies were fixed in 10% buffered formalin less than 30 mins after the resection, for a minimum of 12 hours. The formalin fixed tissue was dehydrated through graded alcohols and embedded in paraffin wax, from which 4 μm thick sections were cut for routine haematoxylin and eosin staining and a panel of immunohistochemical stains. As part of the diagnostic work up, the sections were immunostained for Aβ with immunoperoxidase-labelled antibody 6F3D (DAKO, 1:50), for phosphorylated tau with antibody AT8 (Innogenetics, 1:100), and for this study in addition with antibody against PDGFRβ (RD systems, cat. no. MAB1263, 1:20) to label pericytes. This was performed on a Roche Ventana Discovery automated staining platform following the manufacturer’s guidelines, using biotinylated secondary antibodies and streptavidin-conjugated horseradish peroxidase and diaminobenzidine as the chromogen. The extent of parenchymal Aβ deposition was assessed semi-quantitatively as absent, moderate or severe by a neuropathologist. In addition, to objectively quantify Aβ deposition, the images of the immunoperoxidase label for Aβ were imported into ImageJ, and split into red, green and blue channels. Then, the light intensity in the blue channel (which gave best distinction of the immunoperoxidase label from the background tissue haemotoxylin labelling) was measured in the region of the biopsy where diameters were measured, normalised by the intensity in a region of the section showing no visible Aβ label and converted to a percentage of light absorbed by the Aβ. Normalising by the intensity in a (tissue-free) region without any tissue absorption gave values that were 5.8±0.5% larger, which did not materially change the form of the graphs.

The mean age of patients without Aβ deposition was 50.5±5.5 (n=6, 4 women and 2 men), and of those with Aβ deposition was 62.1±4.2 (n=7, 4 women and 3 men, not significantly different, p=0.11). Regressing mean capillary diameter against age from all patients, or from the patients lacking Aβ deposition, showed that there was no significant dependence on age (p=0.5 and p=0.82 respectively).

Images were analysed to assess capillary diameter with the analyst blind to the level of Aβ and tau deposits (viewing only the PDGFRβ channel). A standard 5×4 grid of 20 squares (each with sides 400 μm long) was superimposed on each image, and all capillaries with clearly demarcated endothelial walls visible in each square had their diameter measured. The image squares were treated as the experimental unit for statistical analysis. Analysis of the diameter as a function of distance from the nearest visible pericyte employed a subset of all the measured diameters, because often no pericyte was visible on some short capillary segments.

### Immunohistochemistry of non-biopsy tissue

Human and rat brain slices were fixed in 4% paraformaldehyde for 1 hour, washed 3 times in phosphate-buffered saline (PBS), then blocked in goat serum. Primary antibodies for PDGFRβ (Santa Cruz, cat. no. sc432, 1:200) or α-SMA (Santa Cruz, cat. no. CGA7, 1:200) were applied overnight, followed (after washing in PBS) by application overnight of Alexa Fluor 647 conjugated secondary antibodies (ThermoFisher, cat. no. A-21245, 1:200). Slices were then washed once in PBS containing DAPI nuclear stain (1:50,000) for 10 mins and then washed again in PBS. After cover-slipping, slices were imaged on a Zeiss LSM700 confocal microscope.

### Statistics

Data normality was assessed with Shapiro-Wilk tests. Comparisons of normally distributed data were made using 2-tailed Student’s t-tests. Equality of variance was assessed with an F test, and heteroscedastic t-tests were used if needed. Data that were not normally distributed were analysed with Mann-Whitney tests. P values were corrected for multiple comparisons using a procedure equivalent to the Holm-Bonferroni method (for N comparisons, the most significant p value is multiplied by N, the 2nd most significant by N-1, the 3rd most significant by N-2, etc.; corrected p values are significant if they are less than 0.05). Assessment of whether the slope of linear regressions differed significantly from zero was obtained using the t-statistic for the slope. P values comparing vessel diameters in the absence and presence of drugs were calculated for the last data point in each graph shown, or for an exposure time of 45-60 mins if no graph is shown. An estimate of the sample size needed for a typical experiment is as follows: For a control response of 100%, a response standard deviation of 10%, a response in a drug of 70% (30% inhibition), a power of 80% and p<0.05, <6 vessels are needed in each of the control and drug groups (www.biomath.info/power/ttest.htm). The exact numbers depend on the drug effect size and standard error of the data.

### Calculation of effect of vessel constriction on flow

We assume that pericytes are regularly spaced on capillaries at an interval of 2L. For flow governed by Poiseuille’s law, the resistance of a segment of capillary of length L (from a pericyte soma to midway between two pericytes) and radius r_1_ is given by

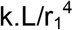

where k is a constant. If Aβ-induced pericyte contraction reduces the capillary diameter from a value of r_1_ at the midpoint between pericytes to r_2_ near the pericyte soma (see Fig. 3a, b, d), then if this reduction is linear with distance the resistance of the capillary segment from the soma to the midpoint is given by

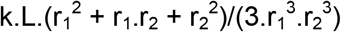

so the factor by which the resistance is altered (compared to that with a uniform diameter r_1_) is

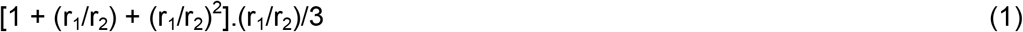

This was used to calculate the predicted flow reduction to be produced by the 30% pericyte constriction reported at pericyte somata in Fig. 3d and the main text. In reality the flow reduction will be greater because the diameter at the pericyte soma is actually larger than at the midpoint between pericytes in control conditions (Fig. 3d), and because Poiseuille’s law does not apply for small capillary diameters for which the effective blood viscosity increases as the diameter decreases below 10 μm^35^.

## Data availability

The data generated in the current study are available from the corresponding author on reasonable request

